# Fatty acid biosynthesis inhibitors fabimycin and triclosan trigger distinct resistance mutations in FabI and potently kill Gram-negative pathogens

**DOI:** 10.64898/2026.04.05.716606

**Authors:** Brett N. Cain, James E. Kent, Marinela Elane, John D. Williams, Myung Ryul Lee, Gee W. Lau, Paul J. Hergenrother, Andrei L. Osterman

## Abstract

In the effort to develop efficacious antibacterials that engage targets for which there is no pre-existing resistance, inhibition of the enoyl-acyl carrier protein reductase FabI has shown promise, with triclosan and fabimycin as representative members of the two major drug classes that show activity against important bacterial pathogens. Here, we use a morbidostat and whole genome sequencing to comprehensively evaluate the resistance profiles that arise in pathogenic bacteria in response to these FabI inhibitors. When assessed against *E. coli*, fabimycin and triclosan were found to induce primarily non-overlapping resistance profiles leading to minimal cross-resistance between the two compounds. Furthermore, *in vivo* evaluation of the prominent resistant mutants indicates poor fitness, with the most fit mutant still susceptible to fabimycin. Collectively, these results suggest the combination use of two antibiotics that engage different positions on the same target as a means to kill pathogenic bacteria and limit resistance.

## INTRODUCTION

Effective antibiotics are a bedrock of modern medicine, but bacteria have adapted, leading to the pandrug-resistant strains now found around the world.^1^ This problem led to more than one million global deaths attributed to bacterial resistance in 2021 alone,^2^ making the development of novel antibiotics and novel antibiotic classes a top priority. Biological targets that are essential and common across pathogenic bacteria, but not mammals, are attractive for antibacterial drug discovery; one such target is the enoyl acyl-carrier protein reductase (FabI), which catalyzes the final, rate-controlling step in bacterial fatty acid biosynthesis (**Figure 1A**).^3^

**Figure 1.**
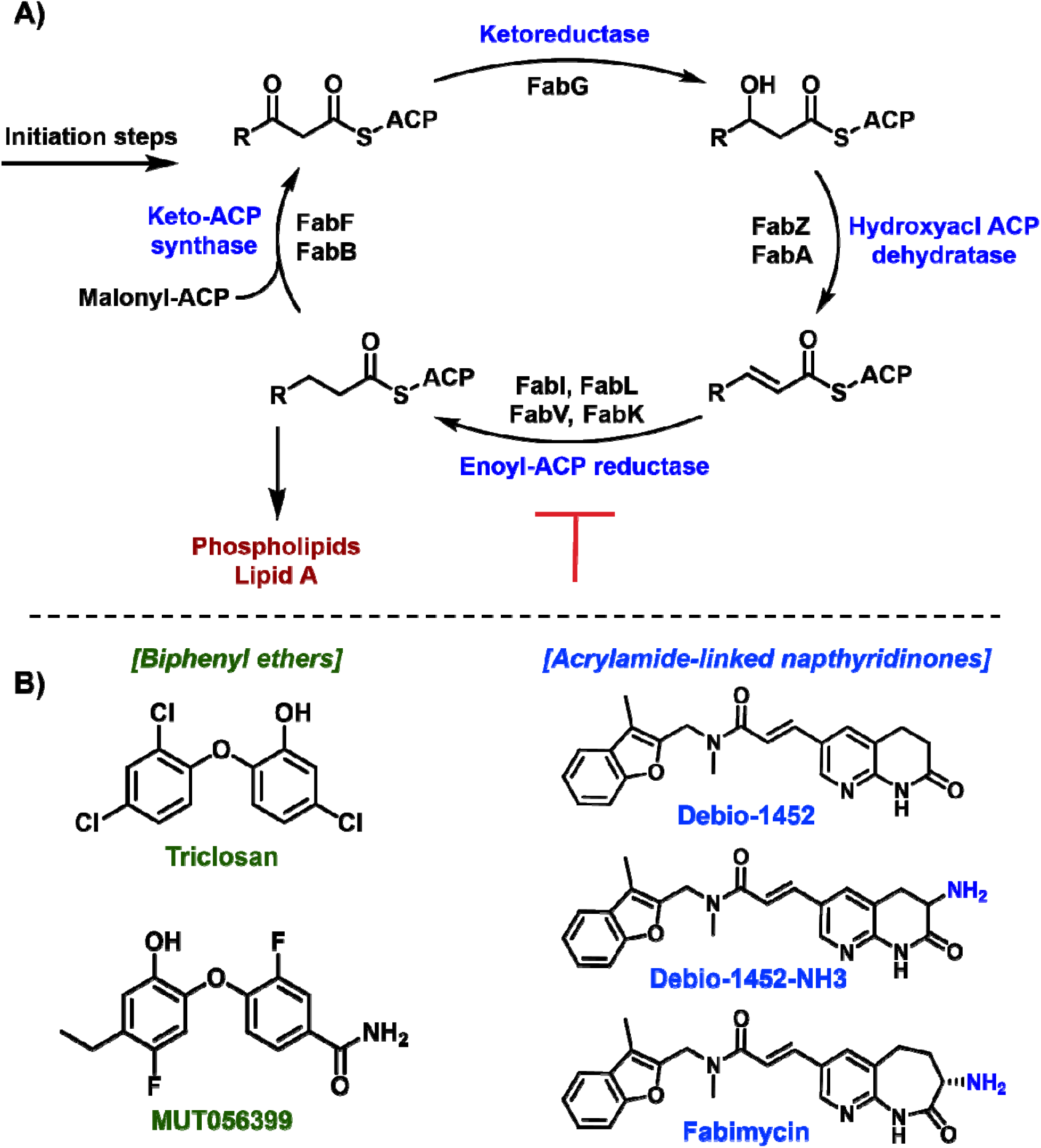
Bacterial fatty acid biosynthesis and important FabI inhibitors. **A)** Fatty acid biosynthesis pathway used by bacteria. **B)** Established small molecule classes of FabI inhibitors.

The most prominent and advanced FabI inhibitors are the biphenyl ethers (represented by triclosan, **Figure 1B**) and the acrylamide-linked napthyridinones (represented by Debio-1452, **Figure 1B**).^4^ Triclosan’s antibacterial activity was first noted in the late 1960’s and it quickly became incorporated into hospital sanitary procedures,^5^ and into many consumer products such as soaps, face washes, hospital scrubs, and even bowling ball finger inserts.^6^ This ubiquity of triclosan stemmed from the belief that it exerted a non-specific killing mechanism similar to other disinfectants (like ethanol), obviating concerns about bacterial resistance. However, it is now recognized that at low concentrations triclosan is a potent inhibitor of bacterial FabI.^7^ Efforts have been made to enhance the bacterial selectivity and potency of triclosan in hopes of generating an antibiotic for systemic use, with MUT056399 being the most advanced derivative (**Figure 1B**).^8, 9^ However, despite these efforts, no antibiotics from the biphenyl ether class have been FDA approved for the treatment of systemic bacterial infections.

The second major class of FabI inhibitors, the acrylamide-linked napthyridinones, arose from early 2000’s target-based screening for inhibitors of *Staphylococcus aureus* FabI.^10^ The first translational candidate in this drug class, Debio-1452 (**Figure 1B**), is interesting as it has a very narrow-spectrum activity for *S. aureus*,^11, 12^ based on differences in the FabI target amongst key Gram-positive bacteria and its inability to accumulate in Gram-negative bacteria. A pro-drug version of Debio-1452 has been assessed in Phase 2 trials for the treatment of infections caused by *S. aureus*^13^ and an analogue with potency against *Neisseria gonorrhoeae* has also been identified.^14^ Recently, using information about chemical traits that facilitate drug accumulation in Gram-negative pathogens, FabI inhibitors Debio-1452-NH3 and fabimycin (**Figure 1B**) were developed that have activity against Gram-negative pathogens including *E. coli, K. pneumoniae, E. cloacae*, and *A. baumannii*, in addition to *S. aureus*.^15, 16^

Antibiotic drug combinations have been developed as a means to increase potency and decrease the ability of bacteria to become resistant, exemplified by the combination of sulfa-drugs/dihydrofolate reductase inhibitors^17, 18^ and the quinupristin/dalfopristin combination^19^. While typical antibiotic combinations target two macromolecules in the same biological pathway, in principle it would be possible to use two compounds that engage different sites on the same essential target. Given the potential of FabI inhibitors, and intrigued by the possibility of developing a combination therapy of two drugs that hit different sites on the same target, here we assess the effect of triclosan and fabimycin in a battery of assays aimed at understanding the mechanism of resistance and translational potential. Provocatively, we show that when exposed to fabimycin or triclosan, *E. coli* evolves distinct sets of mutations in the FabI target, limiting cross-resistance between the two FabI inhibitors.

## RESULTS

### The G148A mutant of FabI is a prevalent and competitive variant that arises in response to fabimycin

Robust understanding of the potential for bacteria to become resistant to an antibiotic is critical for preclinical advancement. The classic methods for these evaluations include assessment of spontaneous resistance frequencies and serial passaging experiments. While these are important workhorse experiments that are operationally simple and have been performed on virtually all classes of antibiotics, a downside is that they assess resistance propensity in a simplistic environment (agar plates or serial passaging in culture) that is not reminiscent of actual human infections. Recently, the development of morbidostat technology has enabled the assessment of bacterial resistance in an adaptive and competitive environment.^20, 21^ The morbidostat is a custom-engineered continuous culturing device in which bacterial mutants evolve through the modulation of drug concentration in response to changing optical densities detected in the device, reflective of bacterial growth. In a typical morbidostat run, bacterial cultures are subjected to continuous growth in six parallel bioreactors over 48-168 hours to capture a wide spectrum of possible mutation events. During the run, the cultures are regularly diluted with drug-free or drug-containing media depending on the observed growth rate/adaptation to gradually increasing drug concentrations until reaching a resistance plateau (See **Supplementary Information** for specific timing and effective drug concentrations). This process, paired with high-coverage whole-genome sequencing (WGS) of the evolving populations, allows for identification of mutants that are not only resistant to the antibiotic, but that are also relatively fit and thus are likely more representative of what might arise in a treatment setting. This methodology has been used to assess the comparative resistomics of several established and experimental antibiotics against prominent pathogens, including *A. baumannii, P. aeruginosa*, and *E. coli*.^20, 22-24^

In the current study, the morbidostat workflow was used to assess *E. coli* ATCC 25922 treated with fabimycin to observe potential evolutionary trajectories towards fabimycin-resistance. Over the course of these experiments numerous *E. coli* variants emerged with missense mutations in the *fabI* gene (**Figure 2, Supplementary Table S1A**). Some of these point mutations had been previously observed using traditional resistance development assays, such as G148A and M159T.^15^ However, the morbidosat-based approach also revealed novel mechanisms of resistance to fabimycin via nine unique point mutations that conferred some level of fabimycin resistance (**Supplementary Table S1A**). A benefit of using the morbidostat to evaluate resistance is the ability to observe competition between the various mutants. Through this lens, the G148A mutant was the most prevalent, outcompeting the other variants (reactors 2 and 6, **Figure 2**) and even competing with long-standing variants once it emerged (reactor 1, **Figure 2**). Other competitive species include H209 variants and M159T, which represent a significant portion of mutants present in reactors 1, 4, and 5 (**Figure 2**).

**Figure 2.**
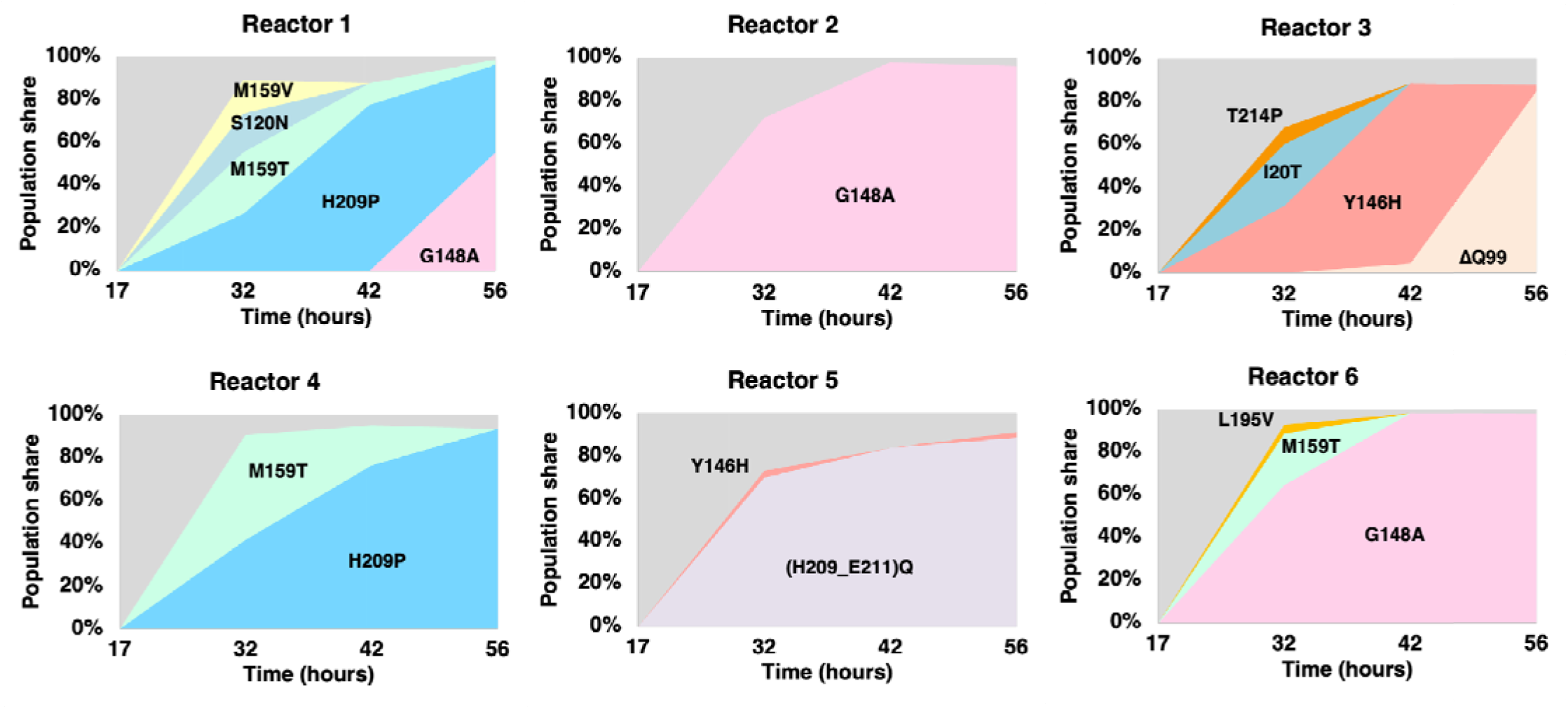
Evolution of *E. coli* in presence of Fabimycin. Mutational profiles of *E. coli* 25922 in response to continuous evolutionary pressure by fabimycin as performed in six parallel reactors (reactors 1 – 6). The X-axis shows four sampling timepoints (hours after the start of the experimental evolution). Colored chart areas reflect relative abundances of respective FabI mutational variants (in percentage of the whole population). In Reactor 5, the event (H209-E211)Q represents the deletion of three residues (H209-E211) and their replacement by a glutamine (Q) residue.

### FabI mutations arising in response to fabimycin or triclosan are largely non-overlapping

In order to compare the resistance profile of fabimycin to another FabI inhibitor, triclosan, the same experimental workflow was repeated, but this time exposing *E. coli* ATCC 25922 to triclosan. Triclosan resistance to *E. coli* BW25113 has been previously explored using the morbidostat^20^, but here we assessed *E. coli* ATCC 25922 to make proper strain-to-strain comparisons. Interestingly, although both fabimycin and triclosan target FabI, the observed mutations in FabI that evolved in response to triclosan pressure were almost completely different than those observed in response to fabimycin (**Figure 3, Supplementary Table S2**). For example, the M159T and G148A variants were two of the most prominent fabimycin-evolved mutants (emerging in 50% of all reactors) whereas these mutants did not often emerge in response to triclosan pressure (M159T observed in 1/6 reactors and G148A in 0/6 reactors). The most prominent variant generated in response to triclosan, A21T, appeared in half of all the reactors. The amino acid residue G93 was found to be mutated to both a valine and serine residue, while not appearing at all under fabimycin drug pressure.The only overlapping mutation found between the fabimycin and triclosan runs was M159T.

**Figure 3.**
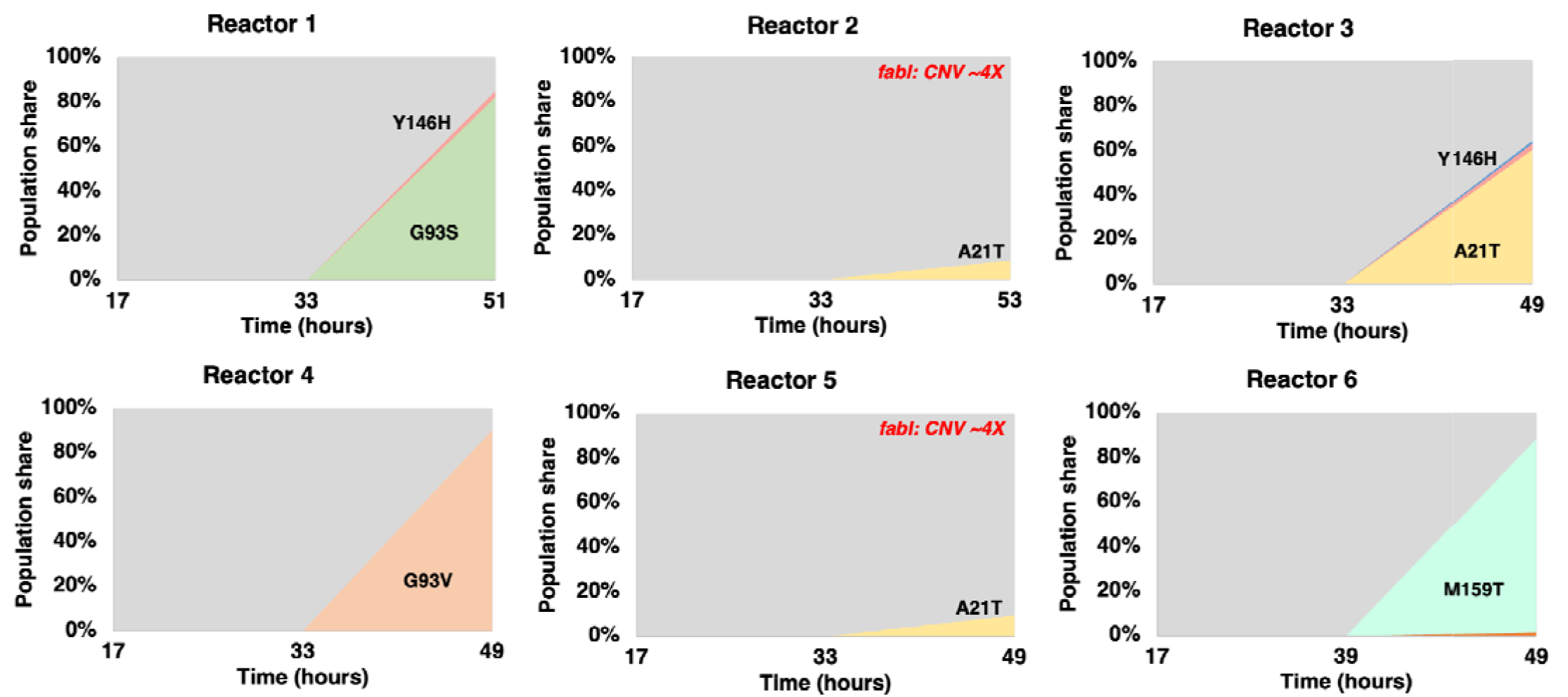
Evolution of *E. coli* in presence of triclosan. Mutational profiles of *E. coli* ATCC 25922 in response to continuous evolutionary pressure by fabimycin as performed in six parallel reactors (reactors 1 – 6). The X-axis shows time after the start of the experimental evolution. Colored chart areas reflect relative abundances of respective FabI mutational variants (in percentage of a population). Red text indicates reactors where amplification of the FabI locus was detected.

With a handful of prominent *E. coli* mutants selected by exposure to fabimycin or triclosan, we sought to understand the rationale and impact of FabI mutations and their influence on the antibiotic activity of the FabI inhibitors. Visualization of point mutations enabled by a co-crystal structure of fabimycin with *E. coli* FabI^15^ revealed that the mutations conferring resistance to fabimycin (shown in blue in **Figure 4**) and triclosan (shown in red in **Figure 4**) largely mapped back to the FabI active site, as expected.

**Figure 4.**
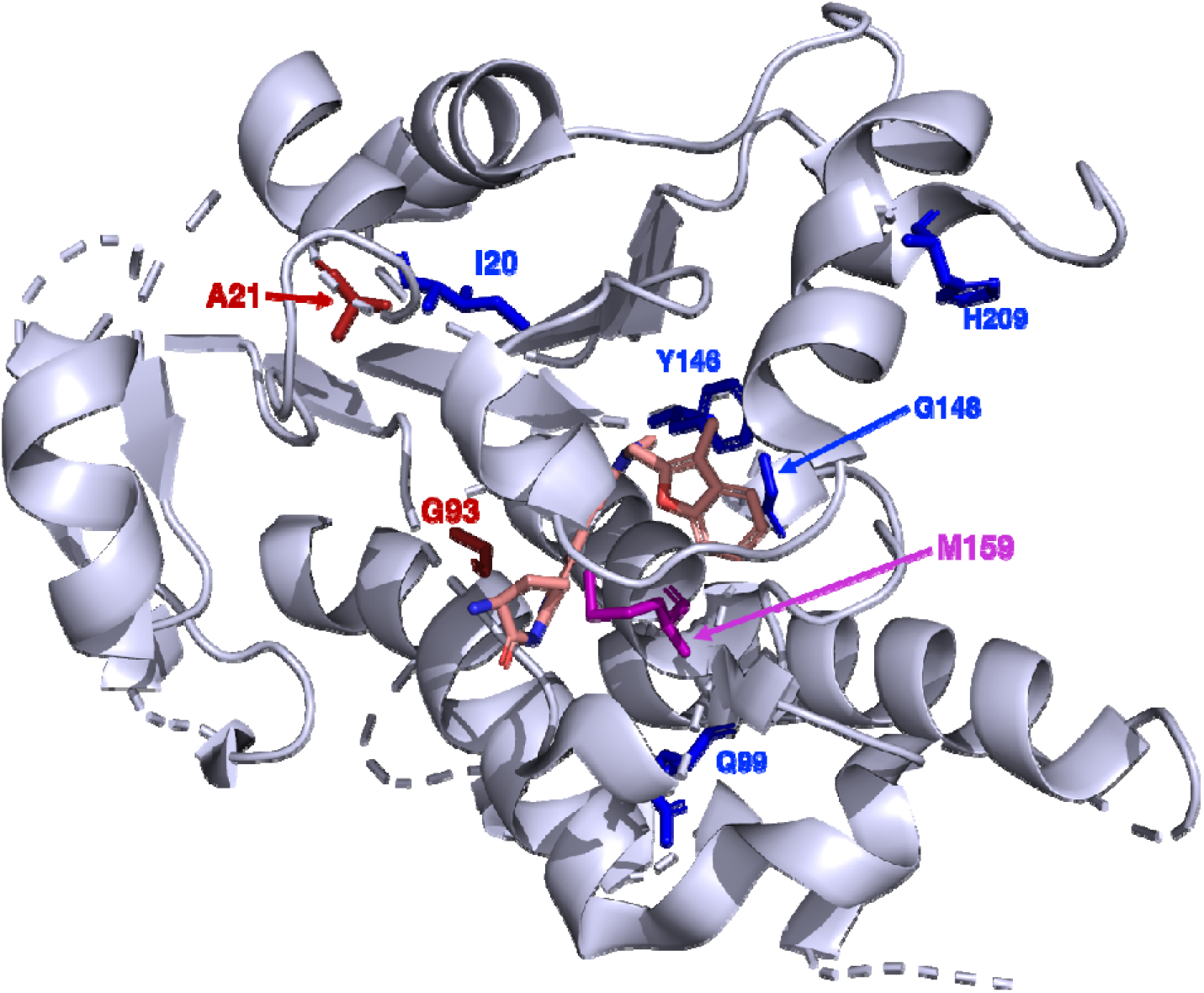
Location of notable FabI mutations. Observed mutated amino acids displayed an X-ray structure of *E. coli* FabI bound to fabimycin (PDB = 7UMW).^15^ Amino acid residues in blue indicate those which emerged under fabimycin pressure, and red indicates those that emerged under triclosan pressure. The sole overlapping mutated residue, M159, is shown in magenta. In this image fabimycin’s carbon atoms are salmon-colored.

### Fabimycin-resistant clones retain sensitivity to triclosan and vice versa

The antibacterial activity of fabimycin and triclosan was evaluated against a panel of clones selected from the morbidostat-based evolution, to assess cross-resistance (**Table 1**). Interestingly, it was fo nd that triclosan retains fair potency versus most of the *E. coli* variants generated in response to fabimycin drug pressure. For example, the competitive and prominent G148A mutant was found to elevate the fabimycin minimum inhibitory concentration (MIC) ∼32-fold, but notably, this mutant remained susceptible to triclosan, with an MIC of 0.62-1.25 μg/mL (a 2-4-fold elevation, **Table 1**). Additionally, the in-frame deletion of Q99, the most impactful mutation on fabimycin activity (ΔQ99; resulting in a 64-fold increase in MIC), did not increase triclosan MICs compared to the wild-type strain, indicating non-overlapping resistance profiles to the two inhibitors (**Table 1** and **Figure 4**). Analogously, *E. coli* harboring the G93V variant of FabI, which arose in response to triclosan challenge and is highly triclosan-resistant (MIC ≥ 20 μg/mL) retains full sensitivity to fabimycin (MIC = 0.62 μg/mL), and this is also true for the G93S mutant with reduced sensitivity to triclosan (MIC = 2.5 μg/mL) and full sensitivity to fabimycin (**Table 1**). In summary, bacteria harboring the vast majority of FabI mutations that arise in response to one inhibitor treatment retain high sensitivity to an alternative Fab Inhibitor (triclosan or fabimycin); the only exceptions are the overlapping M159T mutant, and where the *FabI* gene has been amplified ≥10-fold; these bacteria have reduced sensitivity to both antibiotics (MIC = 5-10 μg/mL, **Table 1**).

**Table 1.** Cross-resistance of clones selected from morbidostat-based evolution of resistance of *E. coli* ATCC 25922 against fabimycin or triclosan. MIC values were measured in triplicate using standard broth microdilution technique utilizing the same growth media as in morbidostat-based experiments. In evolution statistics, “N” reflects the number of reactors (out of 6 total) where a given variant was detected, and “A_max_” reflects the maximal relative abundance (percentage of the population) reached by a given variant.

| FabI variants |  | MIC (µg/mL) |  | Evolution statistics |  |
| --- | --- | --- | --- | --- | --- |
|  |  | Fabimycin | Triclosan | N | A<sub>max</sub> |
| WT |  | 0.62 | 0.31 |  |  |
| Fabimycin-evolved | G148A | 20 | 0.62-1.25 | 3 | 98% |
|  | M159T | 10 | 5 | 3 | 49% |
|  | H209P | 10-20 | 2.5 | 2 | 93% |
|  | (H209-E211)Q | 20 | 1.25 | 1 | 88% |
|  | ΔQ99 | 40 | 0.31-0.62 | 1 | 85% |
|  | Y146H | 10 | 1.25 | 1 | 84% |
|  | I20T | 2.5 | 0.31-0.62 | 1 | 29% |
| Triclosan-evolved | A21T | 1.25 | 1.25 | 3 | 60% |
|  | G93V | 0.62 | ≥20 | 1 | 90% |
|  | M159T | 10 | 5 | 1 | 87% |
|  | G93S | 0.62 | 2.5 | 1 | 82% |
|  | Amplified FabI (10-16X) | 10 | 5 | 1 | 34% |

0.31–1.25 µg/mL
2.5–5 µg/mL
10 µg/mL
20 µg/mL
40 µg/mL

We also assessed a broader panel of *FabI* mutant clones isolated from our previous morbidostat-based experimental evolution study of resistance to triclosan in a model strain of *E. coli* BW25113^20^ or their resistance/susceptibility to fabimycin (**Supplementary Table S1E**). Of the three additional mutational sites (I192, T194 and F203) only the latter was associated with 4-8-fold increase of MIC to fabimycin comparable to 8-16-fold increase of MIC to triclosan. However, no mutations of F203 were detected in this study in the course of experimental evolution under pressure of FabI inhibitors in the strain *E. coli* ATCC 25922.

### FabI inhibitor combination suppresses spontaneous resistance

Intrigued by the mostly non-overlapping point mutations emerging in response to drug exposure and relative lack of cross-resistance between the two FabI inhibitors, experiments were conducted to compare the propensity of bacteria to become resistant to these two antibiotics as sole agents and in combination. Using the large inoculum method, this experiment was executed using *E. coli* BW25113, evaluating frequencies of resistance at 4X-32X the MICs of sole agents (**Figure 5A**). As a single agent against *E. coli*, the frequencies of resistance for triclosan are 10^-9^-10^-10^ and a mutant prevention concentration (MPC) reached at 32X the MIC (2 μg/mL) (**Figure 5A**). Fabimycin’s MPC was also reached at 32X the MIC (16 μg/mL) and with frequencies of resistance comparable and often less frequent compared to triclosan (**Figure 5A**).

**Figure 5.**
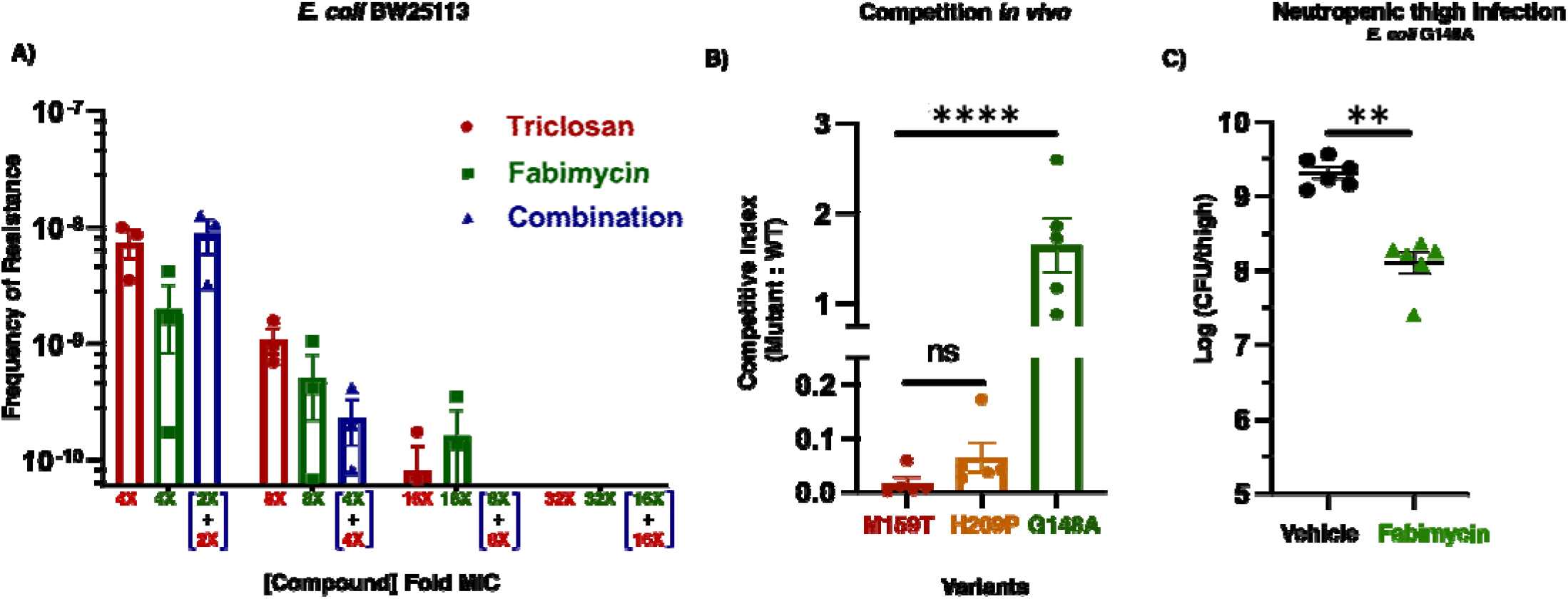
Spontaneous frequency of resistance and analysis of important *E. coli* mutants. **A)** Challenging *E. coli* BW25113 with a range of concentrations of FabI inhibitors and assessing frequency of resistance. Data represent three replicates for each drug treatment with error bars representing the SEM. **B).** *In vivo* competition experiment during bacteremia infection with prominent *E. coli* mutants versus the wild-type parent strain. For each group, 5 immunocompetent mice were inoculated intraperitoneally with an equal amount of the wild-type and variant bacterial strain to induce bacteremia. After 24 hours, the mice were euthanized and selective media was used to determine amounts of each strain present in the infected lung tissue. For reference, the competitive index for the WT strain is equal to 1.0. Statistical significance was determined by one-way ANOVA with Tukey’s multiple comparisons. NS, not significant. \*\*\*\**P* < 0.0001. Error bars represent standard error of the mean. **C)** Neutropenic thigh infection model with prominent *E. coli* mutant G148A. Mice (groups of six) infected with the *E. coli* ATCC 25922 G148A variant (2.2 * 10^5^ CFU per mouse intramuscular in thigh) were treated with vehicle or fabimycin. Fabimycin was formulated in 17% Cremophor EL, 3% SBE-β-CD in H_2_O and administered intramuscularly at 50 mg/kg TID. The bacterial burden was evaluated 24 hours post-infection. The statistical significance was determined by one-way ANOVA with Tukey’s multiple comparisons. \*\**P* = 0.0023. Error bars represent standard error of the mean.

To determine if the non-overlapping mutational profiles of fabimycin and triclosan would result in a diminished propensity for resistance to arise, we assessed frequencies of resistance against a combination of both drugs by *E. coli* BW25113 (**Figure 5A**). To compare frequencies at a given fold-MIC value, we challenged by using the same effective dose of both antibiotics. For example, the resistance frequency at 4X the MIC for a sole agent was compared to the resistance frequency observed when bacteria were challenged with a combination agar plate containing 2X the MIC of each sole agent (2X MIC concentration of triclosan + 2X MIC concentration of fabimycin = 4X effective FabI inhibitor concentration). Using this dosing regimen, frequencies of spontaneous resistance were observed to be lower at concentrations ≥4X compared to the sole inhibitors MICs. Even though lower concentrations of each individual drug were present, this observation demonstrates the power of having both inhibitors present. Notably, no mutants were observed when 8X the MIC of each inhibitor was present (**Figure 5A**).

### FabI mutations that enable *E. coli* to resist triclosan and fabimycin are not fit *in vivo*

To assess the *in vivo* fitness of *E. coli* mutants identified in the morbidostat, we evaluated the most prominent variants for their ability to compete against the parental wild-type strain in immunocompetent mice. Briefly, bacteremia was induced in mice using a 1:1 mixture of *E. coli* ATCC 25922:mutant variants. After 24 hours, the mice were sacrificed and infected lung tissue plated on selective media to assess bacterial counts. Colony-forming unit (CFU) counts of each were assessed and visualized using a ratio of mutant:wild-type. In this experiment, the M159T variant (the only FabI point mutant that enables resistance to both triclosan and fabimycin, **Table 1**) was shown to be the least competitive among the variants assessed (**Figure 5B**); the H209P mutant was also deemed unfit through evaluation in the same experiment (**Figure 5B**). While bacteria with the G148A mutation demonstrated comparable fitness to the wild-type, as shown in **Table 1** triclosan retains potent activity versus these bacteria. The fitness of prominent variants *in vitro* was also evaluated, but no significant differences were found (as expected for morbidostat-based selection), contrasting with the *in vivo* results (**Figure S1**).

### *In vivo* efficacy model utilizing fit and prominent fabimycin-induced FabI mutant, G148A

Intrigued by the competitive (both in culture and *in vivo*) and prominent nature of the fabimycin-induced G148A mutant, we aimed to explore if bacteria harboring this *FabI* mutation present a translational liability. An *in vivo* model was performed where neutropenic mice were infected with the G148A mutant bacteria and treated with a modest dosing regimen of fabimycin (**Figure 5C**). After just 24 hours, dosing fabimycin intramuscularly at 50 mg/kg TID was effective in reducing the bacterial load over 10-fold despite the elevated *in-vitro* MIC of 20 μg/mL (**Figure 5C** and **Table 1**), suggesting that infections from bacteria harboring this mutation could be treated by fabimycin. Of course, given the sensitivity of this mutant to triclosan (**Table 1**), such bacteria would also be expected to be efficiently treated with a triclosan derivative suitable for *in vivo* use.

### Experimental evolution of fabimycin resistance in *A. baumannii* and *S. aureus*

The morbidostat technology was also used to evaluate fabimycin against another Gram-negative pathogen, *A. baumannii*. This experiment revealed point mutations in FabI, with some variants (like A139V) appearing in half of all the reactions with relatively high abundance (**Supplementary Table S1C**). Interestingly, it was found that many variants survived fabimycin pressure by mechanisms of resistance other than FabI point mutation, such as up-regulation of FabB and efflux pumps; these results highlighting the benefits of using the morbidostat to evaluate resistance, as prior studies did not identify these alternative resistance determinants (**Figure S2 and Supplementary Table S1F**). When fabimycin was evaluated in the morbidostat against another pathogen, Gram-positive *S. aureus*, there were relatively few missense mutations observed in *FabI* unlike the morbidostat runs with Gram-negative species like *E. coli* and *A. baumannii* (**Supplementary Table S1D**). In fact, from all six reactors, only four different missense mutations were observed, which were all detected in low abundance (**Supplementary Table S1D and S1G**). Instead, it was more common for *S. aureus* to become resistant via regulatory mutations upstream of *FabI* and *fabI* locus amplification (**Figure S3**). Altogether, these studies show species-specific pathways to resistance from fabimycin, with *E. coli* largely responding with alterations leading to point mutations in FabI, whereas *A. baumannii* and *S. aureus* leveraging alternative resistance-mediating mechanisms.

## CONCLUSIONS

Relative to typical experiments assessing antibiotic resistance, the morbidostat workflow enables a larger view of genomic changes and includes a built-in assessment of the relative ability of the various arising mutations to compete with one another. This method can also uncover resistance-mediating mutations not detected through traditional agar-based assays, as was true herein for fabimycin.

The success of antibiotics like β-lactams and fluoroquinolones is attributed to their ability to effectively inhibit more than one bacterial target.^25^ Significantly less explored is the concept of drugging a single target with two distinct chemical agents. As demonstrated herein with the dual inhibition of FabI, such a strategy can enable potent bacterial killing and suppress the emergence of resistance. FabI is an excellent candidate for this strategy as it is critical for bacterial survival, and inhibition leads to efficacy *in vivo* as exemplified by the acrylamide-linked naphthyridinone class of compounds like afabicin, Debio 1453, and fabimycin.^13-15^ This antibiotic class has exquisite selectivity for bacterial FabI, has minimal impact on commensal bacteria,^26^ and has activity versus priority pathogens. Additionally, this target is not present in mammalian cells, and has been validated in another context where isoniazid (which inhibits InhA, the mycobacterial version of FabI) is routinely included in treatment regimens to treat tuberculosis.^3^

Essential to this combination strategy are distinctly-arising resistance profiles from each antibiotic, in which resistance-conferring mutations to one drug do not impact the antibacterial activity of the other, a trait shown here with fabimycin and triclosan. Indeed, the most fit and prominent fabimycin-evolved variant G148A is easily inhibited by the action of triclosan, highlighting the complementary nature of these two FabI inhibitors. While triclosan displays toxicity when systemically distributed in mammalian systems^9, 27^ and as such is likely unsuitable for direct translational advancement, some more promising analogues have developed.^9^ The work herein suggests combination evaluation of advanced candidates from both compound classes for their ability to treat bacterial infections and suppress resistance, and provides impetus for development of triclosan derivatives suitable for *in vivo* use. More generally, these results suggest the significant potential of an antibiotic strategy employing inhibitor pairs that have distinct binding sites on the same target.

## Supporting Information

Supporting figures and tables, materials and methods, and experimental procedures (PDF).

## Materials and methods

### Bacterial strains and media

Bacterial strains used in this study, *Escherichia coli* ATCC 25922, *Acinetobacter baumannii* ATCC 17978 and *Staphylococcus aureus* ATCC 29213, were obtained from the American Type Culture Collection (ATCC). A cation-adjusted Mueller-Hinton broth (CA-MHB) with 2% dimethyl sulfoxide (DMSO) was used as a base media for the experimental evolution in the morbidostat and MIC measurements. For morbidostat runs, the autoclaved antifoam SE-15 (Sigma) was added to a final dilution of 1/2,500.

### Morbidostat setup and programming

Our implementation of the morbidostat approach has been described in our previous studies with GP-6, ciprofloxacin and triclosan.^20, 23 28^ Briefly, we use a custom-engineered morbidostat device, which is a computer-controlled bundle of 6continuous culturing bioreactors where the culture density (measured by OD_600_) is controlled by varying an antibiotic concertation in media. Automated regular dilutions (20% by volume, every 15 – 30 min) are performed using a controlled mixture of media from two feed bottles: (i) one, containing drug-free media and (ii) another one, with drug-containing media. The composition of mixture is defined by the algorithm enabling a gradually growing selective pressure driving the evolution of higher drug resistance. The detailed description of morbidostat implementation, program logics and parameters of automated dilutions are provided on GitHub (https://github.com/sleyn/morbidostat_construction).

### Evolution of fabimycin resistance in *E. coli* ATCC 25922

The morbidostat run of *E. coli* ATCC 25922 with fabimycin (FBM) was performed, and the results were analyzed as previously described.^20, 22, 23^ The workflow included the following major steps. First, 6 log-phase starter cultures (OD_600_ = 0.02) prepared from glycerol stocks originating from 6 individual colonies were used to inoculate 6 parallel bioreactors of a custom-engineered morbidostat instrument and subjected to continuous growth over 56 hrs. During the run, the cultures were regularly diluted with drug-free or drug-containing media depending on the observed growth rate/adaptation to gradually increasing FBM concentration up to 10 mg/L (within the first 40 hrs), and then to 50 mg/L. Genomic DNAs from 24 samples collected at time points 17 hrs, 32 hrs, 42 hrs and 56 hrs were a subject of Illumina WGS at 300-600x genomic coverage. The sequencing data were mapped to reference genomes to identify single-nucleotide variants (SNVs), short indels, and copy number variants (CNVs). IS element insertion events were mapped using the specialized software iJUMP.^22^ Mapping and assessment of relative abundance of all mutational events was performed using the established computational pipeline as previously described^22, 29^ and briefly outlined below (“Sequencing data analysis, variant calling and ranking”). Upon identification and ranking of mutational events in potential driver genes (**Supplementary Table S1A**) by a combination of criteria (as described below), we have isolated from the retained samples and analyzed a panel of representative clones. Mutations were mapped by WGS and MIC values for fabimycin (MIC^FBM^) and triclosan (MIC^TCS^) were determined by broth microdilution method for a non-redundant set of 7 clones (**Supplementary Table S1E)**.

### Evolution of triclosan resistance in *E. coli* ATCC 25922

The morbidostat run of *E. coli* ATCC 25922 with triclosan (TCS) was performed, and the results were analyzed as described above. The same starter cultures were used to inoculate 6 morbidostat bioreactors and subjected to continuous outgrowth for 55 hrs. During the run, the TCS concentration was gradually increasing up to 5 mg/L (within the first 23 hrs), and then to 20 mg/L (limit of TCS solubility). Genomic DNAs from 18 samples collected at time points 17 hrs, 33 hrs, and 49 - 55 hrs were a subject of Illumina WGS at 400-800x genomic coverage. Upon identification and ranking of mutational events (**Supplementary Table S1B**), we have isolated and analyzed a panel of representative clones by WGS. MIC values for triclosan (MIC^TCS^) and fabimycin (MIC^FBM^) were determined for a non-redundant set of 6 clones (**Supplementary Table S1E)**.

### Evolution of fabimycin resistance in *A. baumannii* ATCC 17978

The morbidostat run of *A. baumannii* ATCC 17978 with FBM was performed, and the results were analyzed as described above. 6 starter cultures (from the same glycerol stocks as in previous studies^29^ )were used to inoculate 6 morbidostat bioreactors and subjected to continuous outgrowth for 64 hrs. During the run, the FBM concentration was gradually increasing up to 25 μg/mL (∼10x of WT-MIC, within the first 53 hrs), and then to 125 μg/mL (∼ 50x of WT-MIC). Genomic DNAs from 24 samples collected at time points 21 hrs, 41 hrs, 54 hrs and 64 hrs, were a subject of Illumina WGS at 800-1,500x genomic coverage. Upon identification and ranking of mutational events (**Supplementary Table S1C**) we have isolated and analyzed a panel of representative clones by WGS. MIC values for fabimycin (MIC^FBM^) were determined for a non-redundant set of 8 clones (**Supplementary Table S1F)**.

### Evolution of fabimycin resistance in *S. aureus* ATCC 29213

The morbidostat run of *S. aureus* ATCC 29213 with FBM was performed, and the results were analyzed as described above. Starter cultures prepared from glycerol stocks originating from 6 individual colonies were the subject of continuous growth in 6 morbidostat bioreactors and subjected to continuous outgrowth for 60 hrs. During the run, the FBM concentration was gradually increasing up to 0.03125 μg/mL (∼5x of WT-MIC). Under drug pressure, cultures showed a tendency to produce a biofilm interfering with monitoring OD600 and thus preventing us from a more significant increase in drug concentration. This tendency was particularly strong in reactors #1 and #3 that were terminated earlier (at 30 hrs). Genomic DNAs from 20 samples collected at time points 20 hrs, 29 hrs, 44 hrs and 60 hrs, were a subject of Illumina WGS at 300-600x genomic coverage. Upon identification and ranking of mutational events (**Supplementary Table S1D**), we have isolated and analyzed a panel of representative clones by WGS. MIC values for fabimycin (MIC^FBM^) were determined for a non-redundant set of 7 clones (**Supplementary Table S1G)**.

### Sequencing data analysis, variant calling and ranking

Genomic DNA extraction, WGS library preparation, sequencing, and raw reads processing were as previously described ^23, 28^. Briefly, upon adapter and quality trimming (BBDuk from BBTools suite v. 38.42, https://sourceforge.net/projects/bbmap/), the reads were aligned to reference genomes (BWA MEM v0.7.17 ^30^). LoFreq Viterbi module was used to refine alignment near indel regions ^31^. Base Quality Score Recalibration was made by Genome Analysis Toolkit (GATK) modules BaseRecalibrator and ApplyBQSR ^32^. Sites with high frequency variants were masked from BaseRecalibrator by calling variants in the alignment down-sampled to ∼ 50x coverage with Picard tools v.2.2.1 (https://broadinstitute.github.io/picard/) based on the coverage estimates by mosdepth v. 0.2.6 ^33^. The VCF file for BaseRecalibrator “--known_sites” option was produced by GATK HaplotypeCaller. All other SAM and BAM files manipulations (sorting, indexing, merging and splitting) were performed with Samtools v1.9 ^30^. SNP and indels were identified using LoFreq v2.1.3 ^31^.

Insertion sequence (IS) elements rearrangements were identified by a developed iJump tool (https://github.com/sleyn/ijump). IS elements in the reference genomes were predicted using BLAST against ISFinder database ^34^. Predicted effects of mutations were assigned with SnpEff v4.3 ^35^. VCF files manipulations were performed with bcftools v1.3 ^36^. Copy number variation (CNV) were predicted by CNOGPro package v1.1 for R ^37^. Reference genomes were downloaded from PATRIC database ^38^. WGS data and assembly of all six unevolved clones of *A. baumannii* ATCC 17978 including variations compared to reference genomes were reported in the previous study ^23^.

Ranking of the observed variants as significant was performed based on their statistics across all six reactors as described ^23, 28^. Briefly, we initially prioritize variants reaching high frequency (or maximal abundance, in a population sample weighted on the total site abundance, ***A***_***max***_) in at least one sample across all 6 reactors. Then, for all genes implicated by at least one variant with ***A***_***max***_ ≥10% we consider all variants (non-preexisting, non-synonymous and with ***A***_***max***_≥2%). The second feature for further ranking of the initially prioritized genes, reflects the overall number of independently occurring variants per gene (***N***_***all***_). In contrast to SNVs and IS-inserts, CNVs (large deletions and amplifications) cannot be accurately estimated in population WGS data. Besides, they cannot be unambiguously assigned to a specific gene, as they typically cover from 3 to more than 30 genes. Therefore, the prioritization (and interpretation) of the observed CNV events relies on the comparison with a list of genes implicated by other types of events. Identification of mutations in clones is straightforward. The observed variants typically match those from respective population data, and their calculated abundance is usually >90%.

### Antimicrobial susceptibility tests

MIC values were measured for the unevolved parental clones and selected evolved clones using broth dilution method as described previously^22^, using the same growth media as in morbidostat-based experiments. The fresh colonies were resuspended in freshly prepared CA-MHB medium and inoculated into a series of wells in 96-well plates containing 2-fold increasing concentrations of each compound. Measurements were performed by constant growth or end-point monitoring of OD_600_ in a BioTek ELx808 plate reader at 37 °C.

## Acknowledgements

This work was supported by the University of Illinois and the NIH (R01AI176523 and R01AI167977).

## DATA AVAILABILITY

Clonal and population sequencing data are available in the SRA database by BioProject accession number PRJNA1445470, aside from ‘A’ samples corresponding to the starting points for A. baumannii ATCC 17978, which are available in BioProject accession number PRJNA1016345: reactor 1 - BioSample: SAMN37382541, reactor 2 - BioSample: SAMN37382545, reactor 3 - BioSample: SAMN37382549, reactor 4 - BioSample: SAMN37382553, reactor 5 - BioSample: SAMN37382557, reactor 6 - BioSample: SAMN37382561).The *A. baumannii* ATCC 17978 reference genome is available in the European Nucleotide Archive by sample accession number ERS4228590.

