## Supplemental materials for "Fatty acid biosynthesis inhibitors fabimycin and triclosan trigger distinct resistance mutations in FabI and potently kill Gram-negative pathogens"

**CONTENTS**

**Supplementary Figures S1-S3.**

**Supplementary Tables S1A-S1G available in separate Excel sheet.**

**A.**

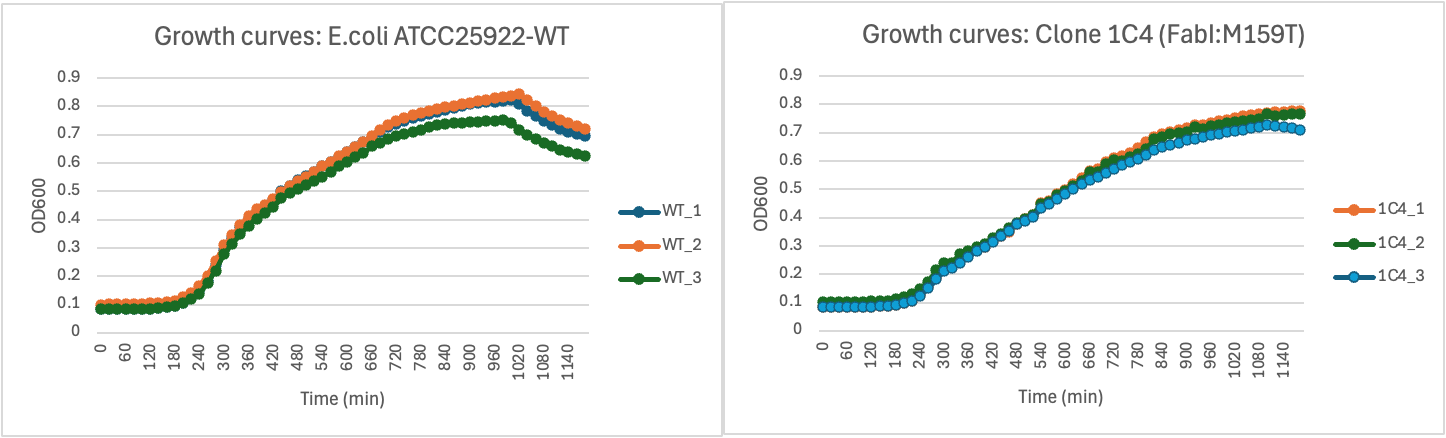

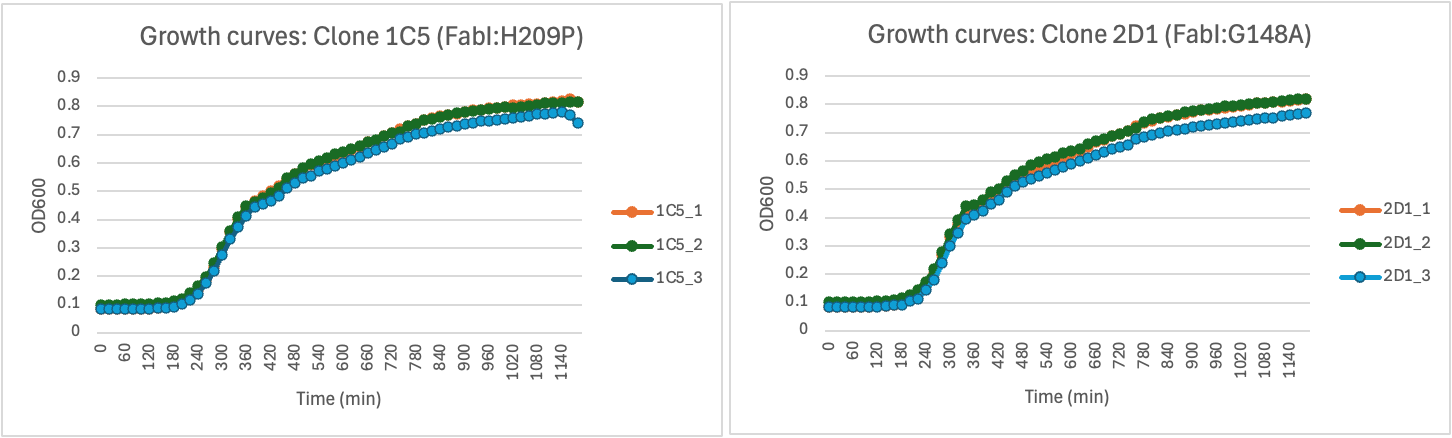

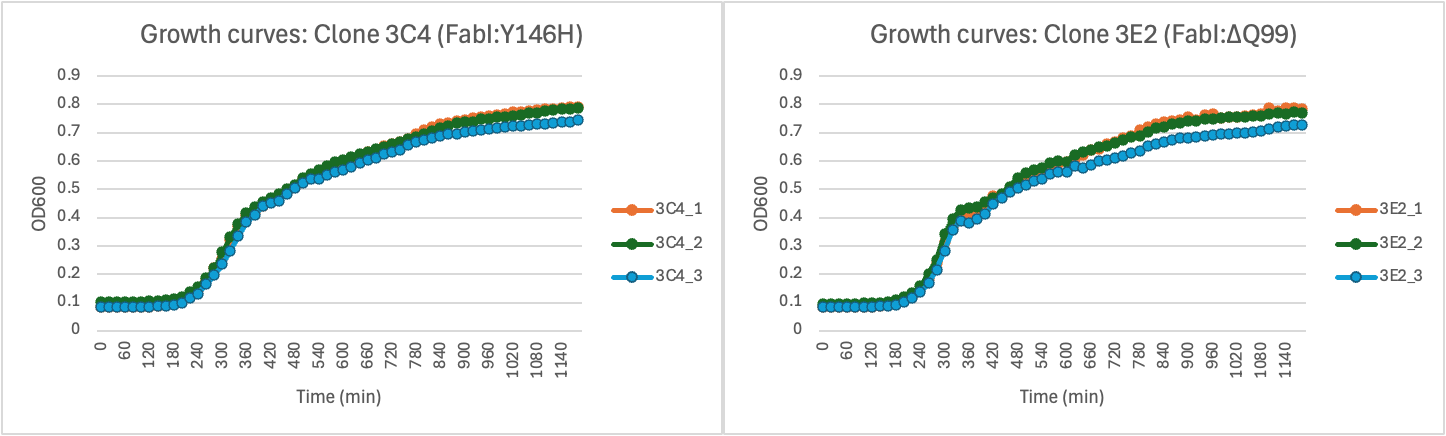

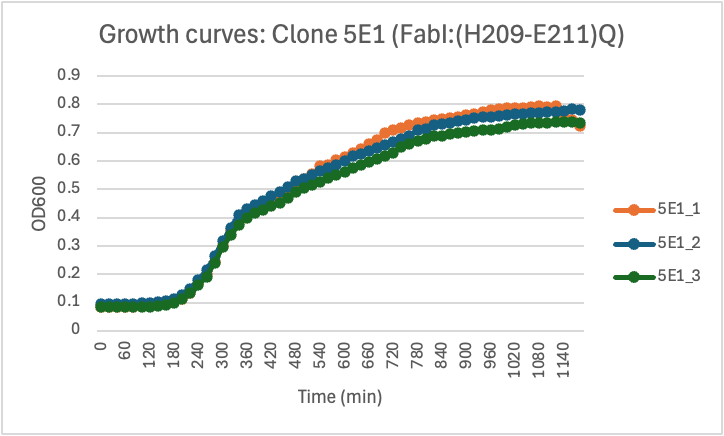

**B.**

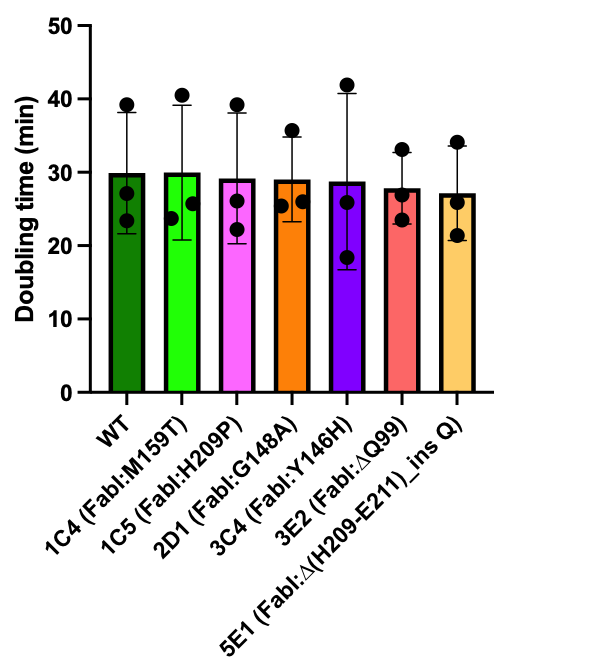

**Supplementary Figure S1**. **Comparing fitness of fabimycin resistant clones and unevolved *E. coli* ATCC 25922 by growth curves.** **A.** Growth curves of unevolved *E. coli* ATCC25922 (**WT**, MIC^FBM^= 0.625 μg/mL) and a panel of clones selected and characterized from morbidostat-based experimental evolution of fabimycin resistance in *E. coli* ATCC25922: **1C4** (FabI:M159T, MIC^FBM^= 10 μg/mL); **1C5** (FabI:H209P, MIC^FBM^= 10-20 μg/mL); **2D1** (FabI:G148A, MIC^FBM^= 20 μg/mL); **3C4** (FabI:Y146H, MIC^FBM^= 10 μg/mL); **3E2** (FabI:∆Q99, MIC^FBM^= 40 μg/mL); **5E1**(FabI:∆(H209-E211)_ins Q, MIC^FBM^= 20 μg/mL). Growth curves were obtained by propagating respective inoculates in triplicates in CA-MHB media in 96-well microtiter plates over 20 hrs at 37^o^C monitoring growth by OD_600_ measurements in a BioTek ELx808 plate reader. **B.** Doubling times were calculated from growth curves using the online tool (<http://dashing-growth-curves.ethz.ch/>) from: Reiter, M.A., Vorholt, J.A. Dashing Growth Curves: a web application for rapid and interactive analysis of microbial growth curves. BMC Bioinformatics 25, 67 (2024). <https://doi.org/10.1186/s12859-024-05692-y>. No statistically significant differences were observed in binary comparisons of doubling times between WT and clones (by one-way ANOVA in GraphPad Prism 10).

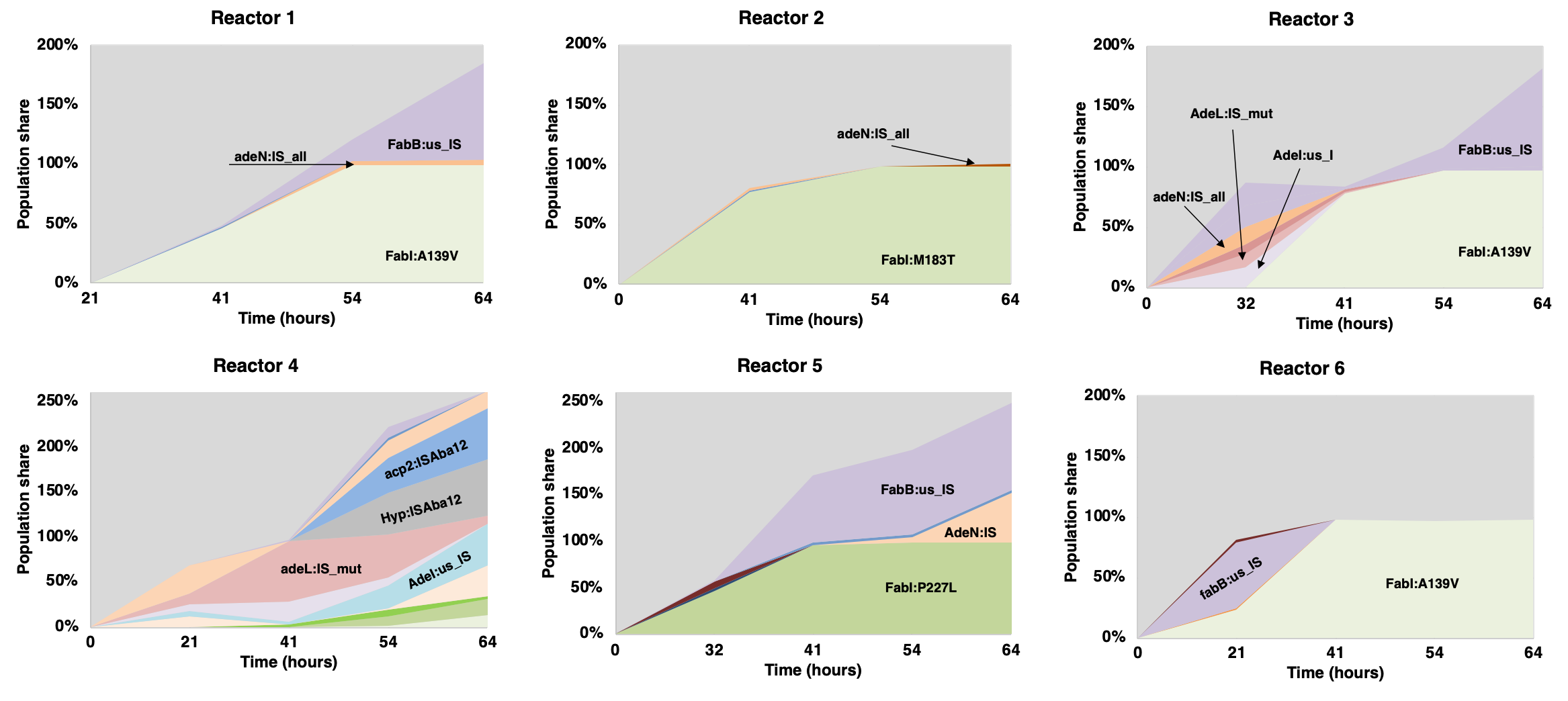

**Figure S2. Morbidstat run of fabimycin versus *A. baumannii* ATCC 19798**. Mutational profiles of *A. baumannii* 19798 in response to continuous evolutionary pressure by fabimycin as performed in six parallel reactors (reactors 1 – 6). The X-axis shows time after the start of the experimental evolution. Colored chart areas reflect relative abundances of respective mutational variants (in percentage of a population).

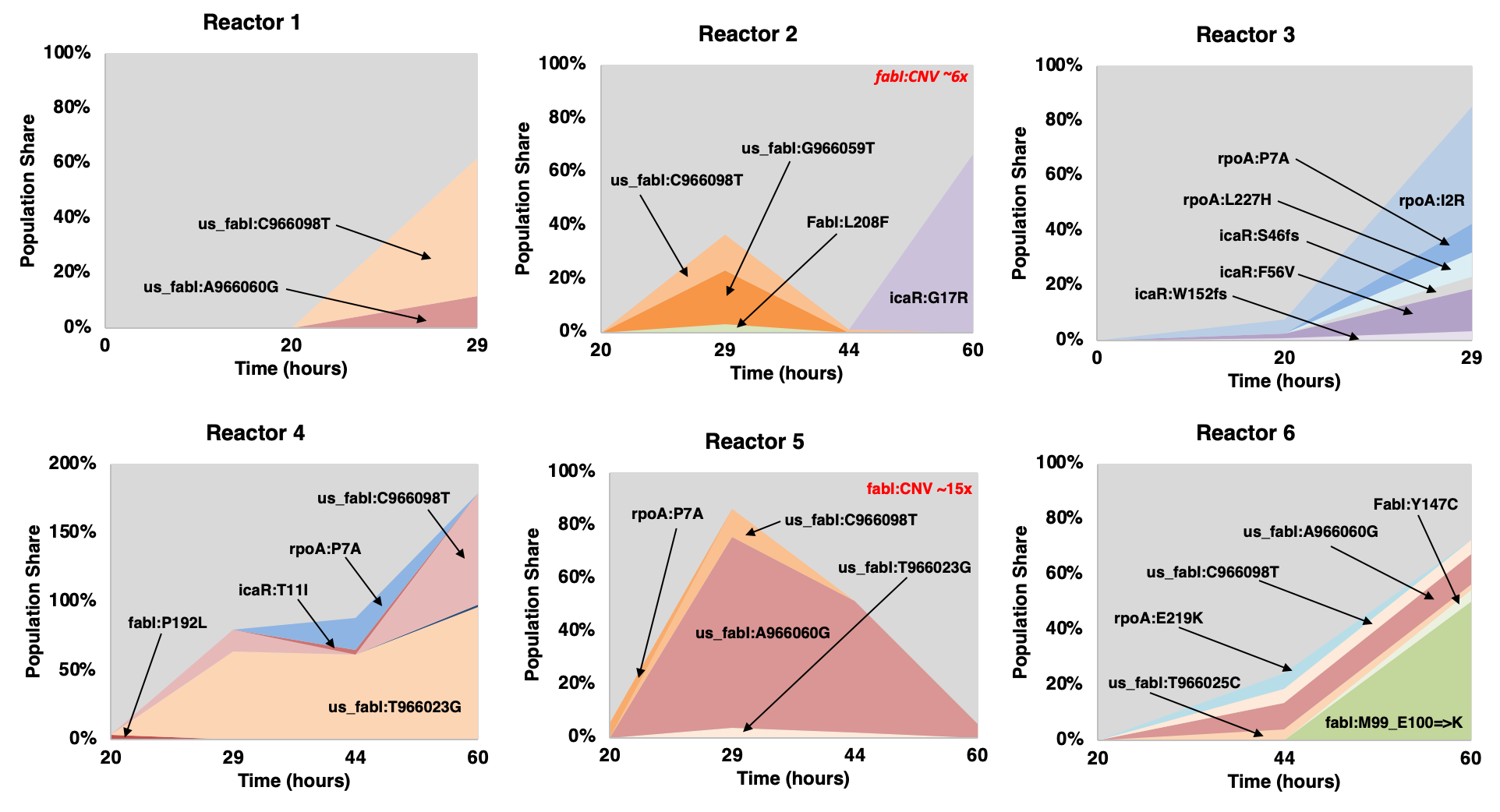

**Figure S3. Morbidostat runs of fabimycin versus *S. aureus* ATCC 29213.** Mutational profiles of *S. aureus* 29213 in response to continuous evolutionary pressure by fabimycin as performed in six parallel reactors (reactors 1 – 6). The X-axis shows time after the start of the experimental evolution. Colored chart areas reflect relative abundances of respective mutational variants (in percentage of a population). Red text indicates reactors where amplification of the FabI locus was detected.
